# Fe_3_O_4_ nanoparticles shell amplify charge-extraction efficiency in *Dunaliella* photovoltaics

**DOI:** 10.1101/2024.06.09.598106

**Authors:** Hao-Hong Chen, Jing-xuan Wu, Jian-Guo Jiang

## Abstract

Microbial biophotovoltaics (BPVs) harness photosynthetic microorganisms to convert light energy into electricity, making them highly attractive for renewable energy production. However, current BPVs typically exhibit low power densities, primarily due to inefficient electron transfer processes and the need for close contact and high interfacial area. Here, we propose a novel method of enhancing *Dunaliella*-based BPVs using Fe_3_O_4_ nanoparticle coatings. The Fe_3_O_4_-coated *Dunaliella* cells (DS@Fe_3_O_4_) establish intimate contact with the cellular electron transfer machinery and maximize the interfacial area, significantly improving electron transfer efficiency and reducing internal resistance. This approach achieved higher power outputs compared to native *Dunaliella* BPVs, with an optimal Fe_3_O_4_ concentration of 2 mg/mL yielding the best performance. In contrast, SiO_2_ coatings on Fe_3_O_4_ (Fe_3_O_4_@SiO_2_) reduced electron transfer efficiency. These findings demonstrate that Fe_3_O_4_ nanoparticle coatings provide a superior method for enhancing bio-electrochemical systems, advancing the application of BPVs for sustainable energy solutions and environmental applications.

## Introduction

Microbial biophotovoltaics (BPV), one type microbial fuel cells (MFC)^1–3^, offer a biological solution for renewable energy production by using photosynthetic microorganisms as light absorbers^4–6^. Algae-based BPVs can produce electricity from both light and organic substrates through the electrons generated during algae’s respiration and photosynthesis, which are then transferred to electrodes within the BPV system, positioning them as potential sustainable energy carriers^5,7,8^.

*Dunaliella*, a single-celled microalga, maintains intracellular sodium concentrations across varying external salinities (0.05-5.0 M NaCl) by regulating osmolytes and glycerol metabolism^9^. In bio-electrochemical systems (BES), *Dunaliella* converts light energy into chemical energy and transfers electrons to the electrode under illumination^10,11^. Its culture also serves as the electrolyte, with biomass and salinity concentration benefiting power generation^12–14^. However, challenges in optimizing electron transfer at enhancing bio-material composite stability and functionality are crucial for advancing BPV and BES technologies for broader energy and environmental applications.

Wiring living cells with abiotic interfaces to create electronic biotic/abiotic interfaces is crucial for various bioelectronic applications^15,16^. BES, leveraging bidirectional electron exchange between electroactive microorganisms and conductive surfaces, have emerged as innovative approaches for energy harvesting, resource recovery, and chemical production^17–20^. These systems aim to couple efficient cell catalysis with material catalysts to achieve high thermodynamic efficiency. However, the slow electron transfer at the biotic/abiotic interface significantly limits BES performance. Efforts to improve this electron exchange have focused on enhancing transmembrane electron transfer pathways mediated by membrane-bound redox proteins^21^. Effective electron transfer requires close contact between these proteins and the conductive surface, a challenge with the traditional "top-down" approach where cells attach randomly to prefabricated abiotic surfaces^22^. This method results in inefficient electron transfer due to idle and dead conduits^23^. Recent advances include using nanoparticles to enhance electron transfer, but challenges remain in fully exploiting individual cell capabilities.

Additionally, strategies to improve MFC anodic electrodes, such as increasing bacteria loading capacity with 3D structures and incorporating carbon nanomaterials or metal nanoparticles, have plateaued in enhancing power density^1,24–26^. Overcoming these limitations requires designing anodic electrodes that fundamentally improve charge transfer efficiency to maximize electron extraction from bacterial metabolism. Recent studies have explored using material shells for biological improvements, including cell protection, stability, and biocatalysis, with shell-coating cells with inorganic nanomaterials proving effective^1,27–30^. This approach can enhance microalgal functions, potentially increasing their value as biomass energy sources^31,32^. However, challenges remain in optimizing these bio-material composites to fully exploit microalgae’s potential in BES for addressing energy and environmental issues.

Here, we report on the enhancement of electrochemical performance and electricity generation in *Dunaliella*-based BPVs using Fe_3_O_4_ nanoparticle coatings. Our studies demonstrate that Fe_3_O_4_-coated *Dunaliella* (DS@Fe_3_O_4_) significantly improves electron transfer efficiency and reduces internal resistance, leading to higher power outputs. Additionally, SiO_2_ coatings on Fe_3_O_4_ (Fe_3_O_4_@SiO_2_) reduced electron transfer efficiency, underscoring the importance of material selection. These findings advance bio-electrochemical systems by optimizing bio-material composites for sustainable energy solutions.

## Results

### Formation of DS@Fe_3_O_4_ core-shell structure

We employed a direct modification method to coat Fe_3_O_4_ onto the surface of *Dunaliella* cells in one step. The shell formation was based on the in-situ self-assembly of magnetic nanoparticles on the surface of the algae cells (**Fig. 1a** and **Supplementary Fig. S1**). The formation of the magnetic shell around *Dunaliella* cells was facilitated by the negatively charged surface of the Fe_3_O_4_ material, leading to electrostatic binding with the cells (**Supplementary Fig. S2,** Zeta potential of core-shell Bio-inorganic materials are fully discussed in **Supplementary Note 1**). This interesting phenomenon made us realize that shell formation can occur spontaneously as long as the surface properties of the material and the biological cells match, allowing for a simple modification that minimizes damage to the cells and achieves the desired outcome.

**Figure 1.**
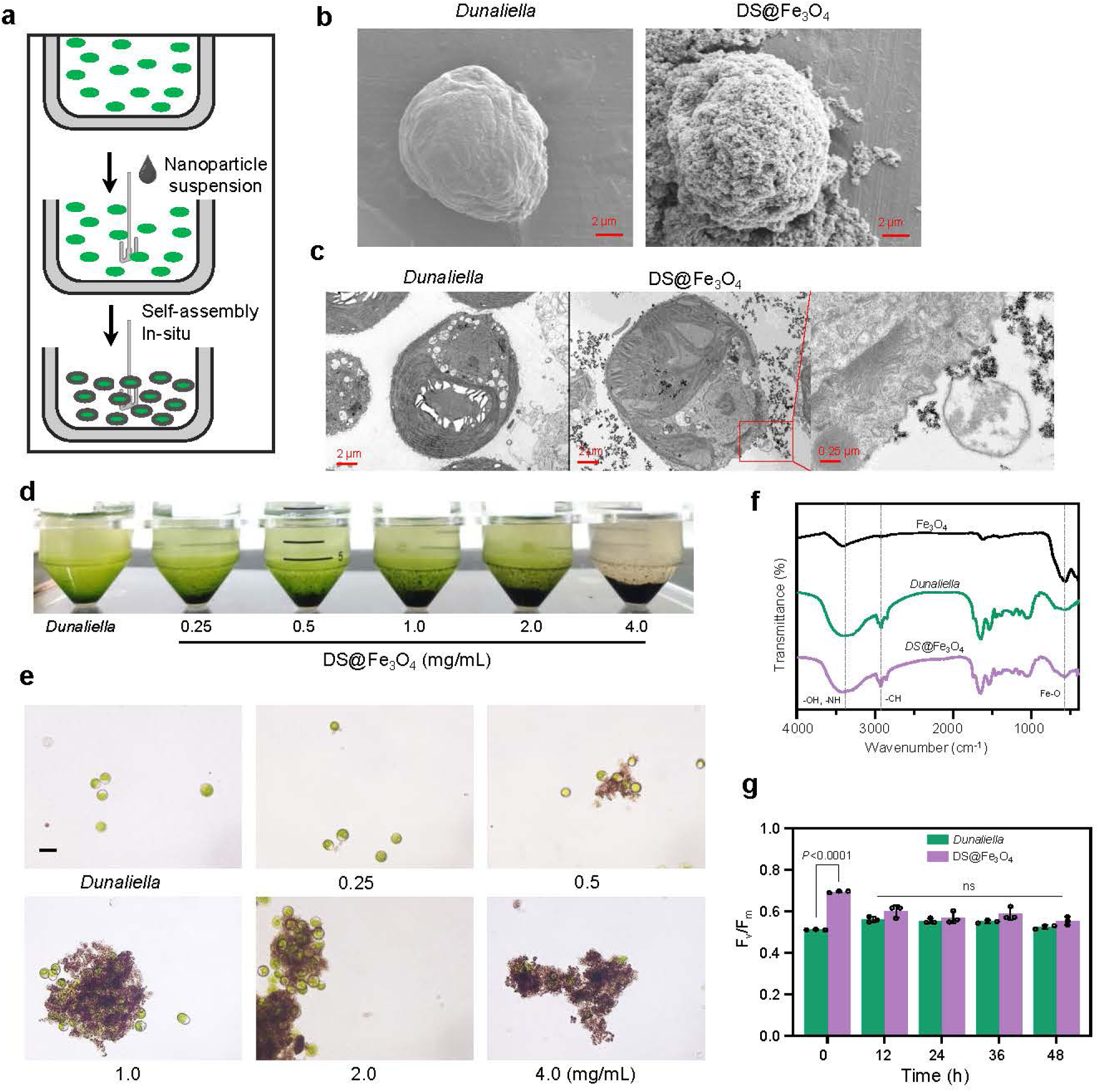
Characterization and analysis of Fe_3_O_4_-coated *Dunaliella* cells. **a.** In-situ self-assembly of Fe_3_O_4_ magnetic nanoparticles on *Dunaliella* surface. **b.** Scanning electron microscope images of native *Dunaliella* cells and cells coated with Fe_3_O_4_ under scanning electron microscope. Scale bar = 2 μm. **c.** Ultrathin section images of *Dunaliella* and DS@Fe_3_O_4_ cells under transmission electron microscope TEM. Scale bar = 2 μm and 0.25 μm (Inset). Algal fluid (**d**) and optical microscope (**e**) of *Dunaliella* cells coated with Fe_3_O_4_ at different concentrations (0, 0.25, 0.5, 1, 2, 4 mg/mL). Scale bar = 10 μm. **f.** Infrared spectra of Fe_3_O_4_, native and Fe_3_O_4_-coated *Dunaliella* cells. **g**. Time-dependent measurements of photosynthetic activity (*Fv*/*Fm*) in the presence of native, Fe_3_O_4_-coated *Dunaliella* cells. Data are presented as mean values ± SD, error bars indicate standard deviations (n =3, biologically independent samples).

Scanning electron microscope (SEM) images and EDX profile revealed that the surface of native *Dunaliella* cells were smooth, with no substances adhering (**Fig. 1b** and **Supplementary Fig. S3**). However, after shell-coating with Fe_3_O_4_, nanoparticles tightly enveloped the surface of the algae cells, making the cell surface rough and granular. The size of native *Dunaliella* was around 10 μm. Following the modification and shell-coating with Fe_3_O_4_, a shell layer 1-2 μm thick formed on the cell surface, resulting in an increased average size of DS@Fe_3_O_4_. In the transmission electron microscopy (TEM) images (**Fig. 1c**), Fe3O4 nanoparticles closely adhered to the surface of the *Dunaliella* cell membrane, but they were only adsorbed on the cell membrane surface and did not enter the cell interior. The internal structure of the *Dunaliella* cells remained intact, possibly because the particle size of Fe3O4 was much larger than the pore size on the cell membrane surface.

Different concentrations of Fe_3_O_4_ nanoparticles were added to *Dunaliella* cell dispersions, setting the concentrations at 0, 0.25, 0.5, 1, 2, and 4 mg/mL. **Fig. 1d** displayed the post-settling appearance of shell-coated *Dunaliella* after 10 minutes, showing increased sedimentation with higher Fe_3_O_4_ concentrations. At 4 mg/mL, the rapid adsorption of Fe_3_O_4_ fully encapsulated the cells, with no green color visible in the supernatant. Optical microscope images in **Fig. 1e** revealed that increasing Fe_3_O_4_ concentrations led *Dunaliella* cells to form larger aggregates. At 0.5 mg/mL, aggregates formed were about 30-40 μm in size; at 1 mg/mL, they were about 80-100 μm; and at 4 mg/mL, a thicker shell of 8-10 μm formed, resulting in over 100 μm irregular aggregates. After Fe_3_O_4_ was electrostatically shell-coated onto the surface of *Dunaliella*, the cells acquired magnetic properties. This disrupted their originally stable negatively charged repulsion system, leading to mutual aggregation and sedimentation, forming clumps.

FTIR analysis was conducted on dried *Dunaliella* cells, DS@Fe_3_O_4_, and nano Fe_3_O_4_ (**Fig. 1f**). By comparing the differences among the three curves, it was observed that DS@Fe_3_O_4_ curve showed a distinct feature from *Dunaliella* curve at 586 cm^-1^, where a strong vibrational absorption peak in Fe_3_O_4_’s infrared spectrum indicates the stretching vibrations of the Fe-O bonds in the inverse spinel structure of Fe_3_O_4_, further confirming that Fe_3_O_4_ had coated the surface of *Dunaliella* cells and grown into a core-shell structure. Both *Dunaliella* and DS@Fe_3_O_4_ samples exhibited spectral features in the amide I region (1600-1700 cm^−1^) at 1650 cm^−1^ and in the amide II region (1500-1600 cm^−1^) at 1540 cm^−1 33^.

Comparing the UV absorption spectra of *Dunaliella* before and after shell-coating, as shown in **Supplementary Fig. S1b**, the nano Fe_3_O_4_ exhibited strong absorption in the 250-400 nm UV range. After shell-coating, the *Dunaliella*’s UV spectrum showed an increase in UV absorption intensity, confirming the successful coverage of Fe_3_O_4_ on the surface of *Dunaliella* cells to form shell-core structure. The full-wavelength absorption spectra curves are consistent in shape, and peaks around 650-700 nm overlap with those of the control group cells at the same concentration. The chlorophyll absorption peak at 680 nm remains unaffected. Therefore, the UV spectroscopy results indicated that shell-coating the cells did not affect the spectral absorption of photosynthetic pigments within the cells.

In summary, we successfully coated Fe_3_O_4_ nanoparticles onto *Dunaliella* cells via in-situ self-assembly, forming a stable core-shell structure. The negatively charged Fe_3_O_4_ bound electrostatically to the algae, creating a 1-2 μm thick shell without damaging the cells. Higher Fe_3_O_4_ concentrations led to larger cell aggregates.

### Maximum quantum yield of Photosystem II and cell growth of DS@Fe_3_O_4_

Photosystem (PS) II is a complex membrane-bound protein located in the thylakoid membranes of oxygenic photosynthetic organisms, responsible for light-driven water oxidation, which produces active electrons (e^-^) and protons^34^. The processes of light capture, excitation transfer, charge separation, and electron transfer within PSII are key initial reactions in photosynthesis, largely determining its overall efficiency^35^. The maximum photochemical efficiency of PSII, *Fv*/*Fm*, represents the intrinsic efficiency of the PSII reaction centers, reflecting the potential photosynthetic capacity of the algae cells^36^.

Photosynthetic microalgae capture sunlight, extract electrons from water, reduce carbon dioxide into sugars, and release oxygen during oxygenic photosynthesis. To understand the ability of algal cells’ PSII to capture solar energy and drive electron transfer activity in the thylakoid membranes, we measured the maximum potential quantum efficiency of PSII in *Dunaliella* cells and DS@Fe_3_O_4_ (**Fig. 1g**). At the initial stages of shell-coating, the DS@Fe_3_O_4_ cells showed an increase in photosynthetic capacity (*Fv/Fm* = 0.69) compared to the *Dunaliella* measurements (*Fv/Fm* = 0.51). This increase may be due to the rapid formation of aggregates after Fe_3_O_4_ coverage, placing the internal algae cells in a hypoxic stress condition that enhances the efficiency of light capture and electron transfer. After 12 hours, the PSII activity began to decline, possibly due to the hypoxic microenvironment within the aggregates, which could suppress the growth conditions of the algae. However, the DS@Fe_3_O_4_ cells retained their original PSII activity for 48 hours, indicating that the aggregation induced by Fe_3_O_4_ almost did not affect the PSII activity of the salt cells.

The inheritance of artificial nano-modifications in subsequent generations of cells remains a challenge in biohybrid system fabrication, as nanoencapsulation of individual cells can inhibit cell growth^15,37^. The growth curves of native *Dunaliella* and DS@Fe_3_O_4_ cells showed that high concentrations (4 mg/mL) of Fe_3_O_4_ inhibited cell division (**Supplementary Fig. S4**). This growth inhibition is advantageous for short-to mid-term applications (several hours to a day) as it provides a stable interface. However, for long-term applications, inevitable cell growth may disrupt the single-cell electron collector and impair electron transfer. Therefore, further development of dynamic/self-repairing nanocoatings or genetically encoded conductive nanocoatings is expected, enabling next-generation intelligent and inheritable single-cell electron collectors for extended applications.

### *Dunaliella* BPV with different cathode electron acceptors

In dual-chamber MFCs, the cathode electron acceptors include ferricyanide^37,38^, permanganate^39,40^, phosphate^41,42^, among others. Ferricyanide can be reduced at the electrode to K_3_[Fe(CN)_6_], and can form a conductive polymer film on the electrode surface, which facilitates electron transfer^37^. Both KMnO_4_ and K_2_S_2_O_8_ possess high redox potentials and have the potential to serve as electron acceptors in MFCs to enhance battery performance^39,41^. Here, the experiment compared the performance of BPVs with K_2_S_2_O_8_ (50 mM), KMnO_4_ (50 mM), and K_3_[Fe(CN)_6_] (50 mM) as cathode electron acceptors, constructing a microbial BPV system with *Dunaliella* as the anodic microorganism (**Fig. 2** and **Supplementary Fig. S5**).

**Figure 2.**
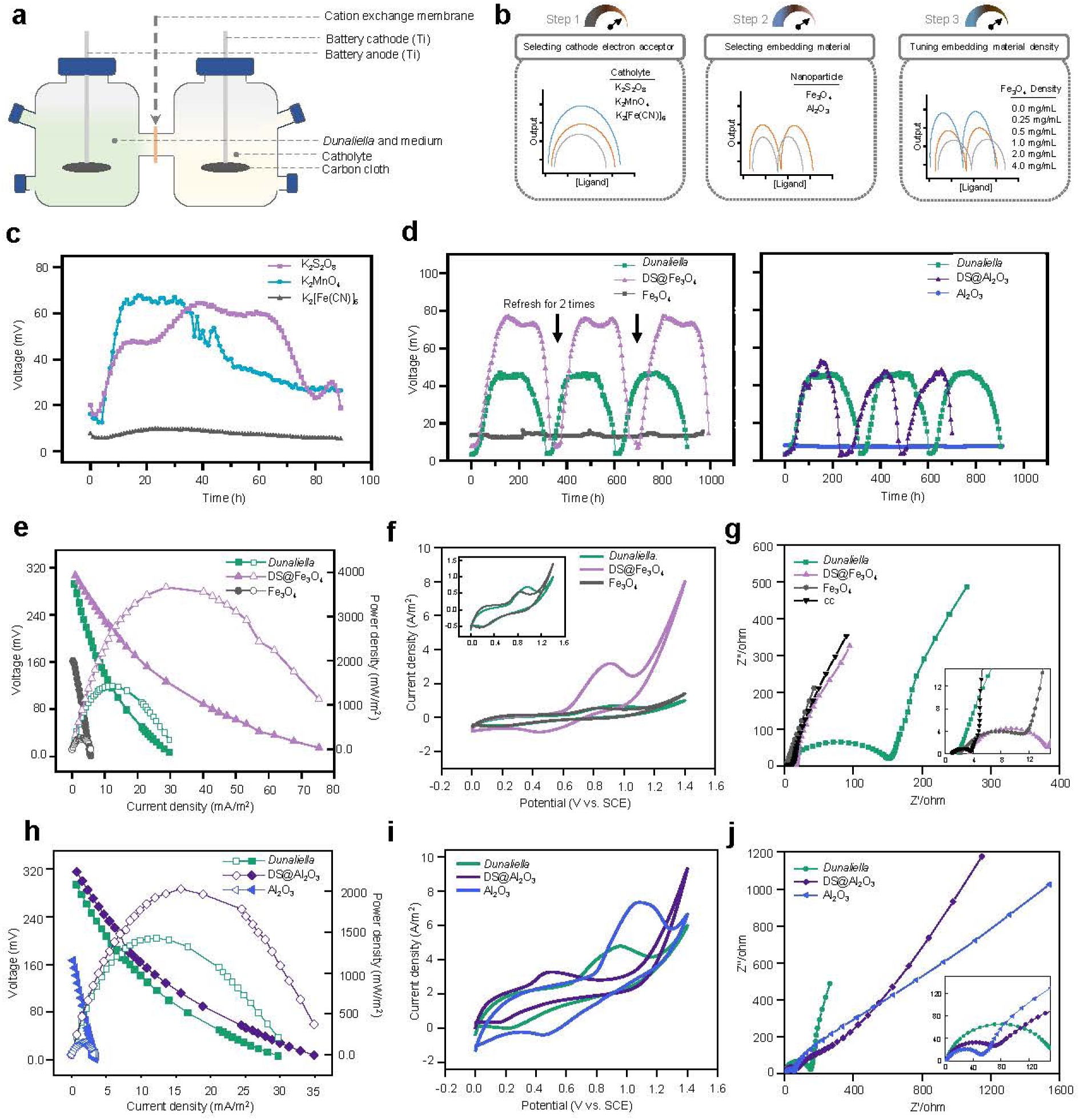
Enhancement and performance evaluation of dual-chamber *Dunaliella* BPVs with Fe_3_O_4_ and Al_2_O_3_ coatings. **a.** Schematic of the dual-chamber *Dunaliella* BPV. Both electrodes of the *Dunaliella* BPVs are made of carbon cloth, connected by titanium wire, forming a closed circuit with the external resistance. **b.** Schematic diagram for the experimental design of *Dunaliella* BPVs. First (Step 1), the catholyte for the BPV was screened from the candidates K_2_S_2_O_8_, K_2_MnO_4_, and K_2_[Fe(CN)_6_] to select the one with the best performance. Next (Step 2), between Fe_3_O_4_ and Al_2_O_3_, the material that was more suitable for shell coating and could efficiently enhance BPVs electron generation performance was compared. Finally (Step 3), a series of concentration gradients for Fe_3_O_4_ were set up to determine the optimal concentration for improving BPVs electron generation efficiency. **c.** Voltage output of dual-chamber BPVs with three different catholytes. The *Dunaliella* cells were set to a final working OD_630_ = 1.2. **d.** Native *Dunaliella*, Fe_3_O_4_, Al_2_O_3_, DS@Al_2_O_3_ and DS@Fe_3_O_4_ BPVs versus time curves for long-term stability and repeated cycling tests. Polarization curves and power output curves (**e**), CV curves (**f**), and EIS spectra (**g**), of native *Dunaliella,* Fe_3_O_4_, and DS@Fe_3_O_4_ BPVs. Polarization curves and power output curves (**h**), CV curves (**i**), and EIS spectra (**j**), of native *Dunaliella,* Al_2_O_3_, and DS@Al_2_O_3_ BPVs. CV curves at a scan rate of 10 mV·s^−1^. CV scans at different scan rates are fully discussed in **Supplementary Note 6**. The *Dunaliella* cells were set to a final working OD_630_ = 1.0.

In the cathode chamber, the electron acceptors on the cathode electrode surface undergo reduction reactions with electrons or protons transferred from the anode, thereby generating current. As shown in the **Fig. 2c**, KMnO_4_ as a cathode electron acceptor achieved higher voltages, but the voltage output was unstable. KMnO_4_, as an electron acceptor, consumes a large amount of H^+^, and to maintain a high electromotive force in the BPVs, acidic solutions such as sulfuric acid must be continuously added to the catholyte to sustain the reaction conditions^40^. Furthermore, a decreasing trend in the battery performance was observed before 40 hours of operation, likely due to the deposition of KMnO_4_ reduction product MnO_2_ on the electrode surface, increasing the battery’s resistance.

The BPV with K_2_S_2_O_8_ as an electron acceptor produced a stable output voltage of 64.76 mV, significantly higher than that of the BPV with K_3_[Fe(CN)_6_] as an electron acceptor (10.67 mV). This was primarily due to the higher standard electrode potential of K_2_S_2_O_8_ in aqueous solution, which was 2.01 V, compared to the lower potentials of permanganate (*E*° =1.70 V) and ferricyanide (*E*° =0.43 V). Based on the experimental results and theoretical analysis, K_2_S_2_O_8_ solution, with its higher redox potential, was selected as the cathode electron acceptor for constructing BPVs.

### Electricity generation characteristics of *Dunaliella* BPV with Fe_3_O_4_- or Al_2_O_3_-coated shell

Fe_3_O_4_ nanoparticles were valued for their biocompatibility, ease of modification, and magnetic separation properties, making them useful in biomedicine and drug diagnostics^43,44^. Peng *et al.* added Fe_3_O_4_ to anodes, improving MFC performance^45,46^. Studies have shown that Al_2_O_3_ improves electrode electrochemical performance, safety, and stability, offering good mechanical strength, electrolyte penetration, and retention, thereby enhancing long-term stability and capacity^47,48^.

Here, a stable core-shell structure was successfully formed by coating Al_2_O_3_ nanoparticles onto *Dunaliella* cells via in-situ self-assembly (**Supplementary note 2** and **Supplementary Fig. S6**). BPVs were constructed using native *Dunaliella*, DS@Fe_3_O_4_, and DS@Al_2_O_3_ as anodic microorganisms, and the resulting voltage-time curves were plotted (**Fig. 2d** and **Supplementary Table S1**). The voltage changes in BPVs are closely related to the growth status of the anodic microorganisms and the extent of microbial enrichment on the anode surface. Based on the voltage trends, the operation of the BPVs could be divided into four phases: startup, rise, equilibrium, and decline. Comparative analysis showed varying equilibrium output voltages for native *Dunaliella*, DS@Fe_3_O_4_, and DS@Al_2_O_3_ BPVs were 47.01 ± 0.29 mV, 77.68 ± 2.19 mV, and 49.64 ± 3.02 mV, respectively, with average operational cycles of 99.3, 106, and 76 hours. Coating Fe_3_O_4_ significantly enhanced the output voltage, demonstrating 65.24% higher than the controls (Native *Dunaliella* and Fe_3_O_4_). These results affirm that current production in BPVs primarily results from the activity of *Dunaliella* and that Fe_3_O_4_ incorporation effectively enhances electron transfer to the electrode, not only reducing startup time and increasing initial voltage but also extending operational duration. Conversely, the highest voltage for the Al_2_O_3_-coated BPV (49.64 ± 3.02 mV) was comparable to that of the *Duanliella* BPVs. Its operational period was significantly shorter, potentially due to Al_2_O_3_’s cytotoxic effects accelerating cell apoptosis, leading to a quicker voltage decline^49^.

Polarization curve and power density curve measurements were taken when the battery’s output voltage reached the equilibrium phase. Before measuring, the battery was left open-circuit for 2 hours. During the measurement, resistance in the circuit was adjusted (from 10,000 Ω to 100 Ω), and the voltage across the battery terminals was recorded (**Fig. 2** and **Supplementary Table S2, 3**). The maximum open-circuit voltages (OCV) for *Dunaliella*, DS@Fe_3_O_4_, and DS@Al_2_O_3_ BPVs were 298.81, 302.85, and 305.16 mV, respectively, and their maximum power densities were 1424.72, 3658.41, and 2030.41 mW/m² (**Fig. 2e**). Comparing polarization curves of BPVs constructed with *Dunaliella*, DS@Fe_3_O_4_, and DS@Al_2_O_3_, the OCVs were similar, indicating that the BPVs’ electricity-generating capacity was comparable before and after shell-coating, with *Dunaliella* being the primary active substance. The apparent internal resistances for the DS@Fe_3_O_4_, DS@Al_2_O_3_, and *Dunaliella* BPVs were 6.68, 12.96, and 17.39 Ω, respectively (**Fig. 2h**). The latter’s resistance was 2.60 and 1.34 times those of the first two, respectively. This decrease in resistance for the DS@Fe_3_O_4_ BPV was due to the electrocatalytic activity and excellent biocompatibility of Fe_3_O_4_ coated, which reduced electron transfer resistance and promotes microbial adhesion on the electrode. Shell-coated algae BPVs showed significantly higher power densities than native *Dunaliella* BPVs due to the enhanced aggregation effect of the shell-coated algae cells under electrostatic action, which aids in microbial enrichment on the electrode surface. Specifically, the power density of the DS@Fe_3_O_4_ (3658.41 mW/m²) and DS@Al_2_O_3_ (2030.41 mW/m²) BPV was 2.57 and 1.8 times that of the *Dunaliella* (1424.72 mW/m²).

In summary, coating *Dunaliella* cells with Fe_3_O_4_ nanoparticles significantly enhanced BPV performance, increasing output voltage by 65.24% and power density to 3658.41 mW/m², due to improved electron transfer and microbial adhesion. Fe3O4’s superior biocompatibility and electrocatalytic properties reduced internal resistance and boosted BPV efficiency.

### Electrochemical performance of *Dunaliella* BPV with Fe_3_O_4_- or Al_2_O_3_-coated shell

Cyclic voltammetry (CV) was applied to reveal the kinetics of redox reactions at the cell-electrode interface in *Dunaliella*, DS@Fe_3_O_4_, and DS@Al_2_O_3_ BPV^4^. The CV analysis provided insights into the electrochemical activity and conductivity of the electrodes based on the positions of the redox peaks, as detailed in **Fig. 2f and 2i**.

The CV curves, characterized by an S-shape typical for electricity-producing microbes, indicated that DS@Fe_3_O_4_ and DS@Al_2_O_3_ exhibited superior biocatalytic activity compared to native *Dunaliella*, as evidenced by the stronger peak currents in the order of DS@Fe_3_O_4_ > DS@Al_2_O_3_ > *Dunaliella*. The biofilm of *Dunaliella* on the carbon felt electrode (CC) exhibited distinct redox waves, with the oxidation peak centered around 0.95 V (vs. SCE), indicating that *Dunaliella* can establish an electron transfer chain on the conductive surface. As for DS@Fe_3_O_4_ BPV, the CV curve exhibited significantly higher oxidation peak currents, centered around 0.91 V (vs. SCE), indicating enhanced electron transfer provided by proteins on the cell membrane. The peak current increased by 4.35 times with Fe_3_O_4_-coated, demonstrating that the redox signals were predominantly due to *Dunaliella*. The complex electrochemical behavior and increased redox currents imply that biocompatible Fe_3_O_4_ acted as an enhancer for long-distance electron transfer efficiency in *Dunaliella*. Both the native *Dunaliella* and the Al_2_O_3_ BPVs exhibited smaller current responses, indicating minimal catalytic activity. Compared to the *Dunaliella* BPV, the DS@Al_2_O_3_ BPV showed 0.75 times increase in oxidation peak current, with the peak centered around 1.09 V (vs. SCE).

CV results demonstrated that BPVs with DS@Fe_3_O_4_ and DS@Al_2_O_3_ as anodes exhibited higher peak currents than *Dunaliella*, significantly surpassing the Fe_3_O_4_ and Al_2_O_3_ controls. This confirmed that the coating of these nanoparticle materials with biological components significantly boosts electron transfer at the electrode surface, making the electrochemical activity of shell-coated *Dunaliella* with Fe_3_O_4_ and Al_2_O_3_ stronger than that of native *Dunaliella*.

We utilized Electrochemical Impedance Spectroscopy (EIS) to analyze the electrochemical activity of DS@Fe_3_O_4_ and DS@Al_2_O_3_ BPVs. Using Zview software along with EIS spectra, we fitted the electrode data using a traditional Randles equivalent circuit model (**Supplementary Fig. S5c**). The EIS diagram showed Nyquist plots consisting of semicircles and a low-frequency straight line (**Fig. 2g** and **2j**).

The electrode impedance data of DS@Fe_3_O_4_ and DS@Al_2_O_3_ derived from the equivalent circuit modeling was provided in **Supplementary Table S4**. The ohmic resistance of the anodes for *Dunaliella,* DS@Fe_3_O_4_, and DS@Al_2_O_3_ was relatively close, ranging between 1 and 2 Ω. The internal resistance of the carbon cloth electrodes in *Dunaliella* and shell-coated *Dunaliella* BPVs were as follows: *Dunaliella* (158.4 Ω), DS@Fe_3_O_4_ (11.13 Ω), and DS@Al_2_O_3_ (50.85 Ω), showing that DS@Fe_3_O_4_ BPV had the lowest bioanode internal resistance, indicating a higher efficiency of electron transfer. The charge transfer resistance (*R*_ct_) for DS@Fe3O4 was 11.13 Ω, significantly lower compared to the native *Dunaliella* bioanode, which was 14.23 times higher. DS@Al2O3 had an *R*_ct_ of 50.85 Ω, which was 3.11 times that of the native *Dunaliella* bioanode. Furthermore, compared to the control Fe_3_O_4_ (no cell) BPV and carbon cloth (CC) electrode, the former showed lower impedance, indicating that Fe_3_O_4_ shell-coating aids in the enrichment of *Dunaliella* cells on the carbon cloth surface and improves biofilm conductivity, enhancing electron transfer capabilities at the anode. This effect was attributed to the conductive nature of the Fe_3_O_4_ shell and the material’s role in assisting algae accumulation on the electrode surface.

Overall, the presence of Al_2_O_3_ and Fe_3_O_4_ reduced the electron transfer resistance between the microorganisms and the anode, enhancing the electrocatalytic activity of the bioanode. The electrodes of the DS@Fe_3_O_4_ BPV exhibited the lowest charge transfer resistance, indicating that Fe_3_O_4_ shell-coating effectively accelerated the electrochemical electron transfer process between the microorganisms and the electrode.

### Electrical generation characteristics of *Dunaliella* BPVs with different Fe_3_O_4_-coated concentrations

To assess the electron transfer capabilities of DS@Fe_3_O_4_ BPVs, we constructed BPVs with different concentrations of Fe_3_O_4_-coated *Dunaliella*, labeled as F0 (0 mg/mL), F1 (0.25 mg/mL), F2 (0.5 mg/mL), F3 (1 mg/mL), F4 (2 mg/mL), and F5 (4 mg/mL).

The output voltage varied across the different concentrations of DS@Fe_3_O_4_ BPVs (**Fig. 3a** and **Supplementary Fig. S7**). The peak voltage outputs for these six groups were in the order of F4 (52.26 mV) > F5 (51.71 mV) > F3 (33.42 mV) > F2 (17.52 mV) > F1 (11.29 mV) > F0 (11.07 mV). F1 and F2 exhibited similar peak output voltages during their equilibrium phases, possibly due to the lower Fe_3_O_4_ concentration (0.25 mg/mL) not being sufficient to coat most of the *Dunaliella*, with many cells remaining suspended in the anode chamber. F4 achieved the highest voltage of 52.26±0.09 mV, which was 4.72 times higher than the native *Dunaliella* BPV (11.07±0.40 mV), with an operation period of 39.4 hours. F5 produced a peak voltage of 51.33 mV, similar to F4, but had the shortest average operation period of 36.7 hours. The operation period tended to decrease for BPVs with Fe_3_O_4_ concentrations greater than 1 mg/mL, likely due to the cytotoxic effects of high Fe_3_O_4_ concentrations or mechanical compression on the algae by excessive Fe_3_O_4_, limiting their viability. These results suggested that BPVs with 2 mg/mL Fe3O4-coated *Dunaliella* were optimal.

**Figure 3.**
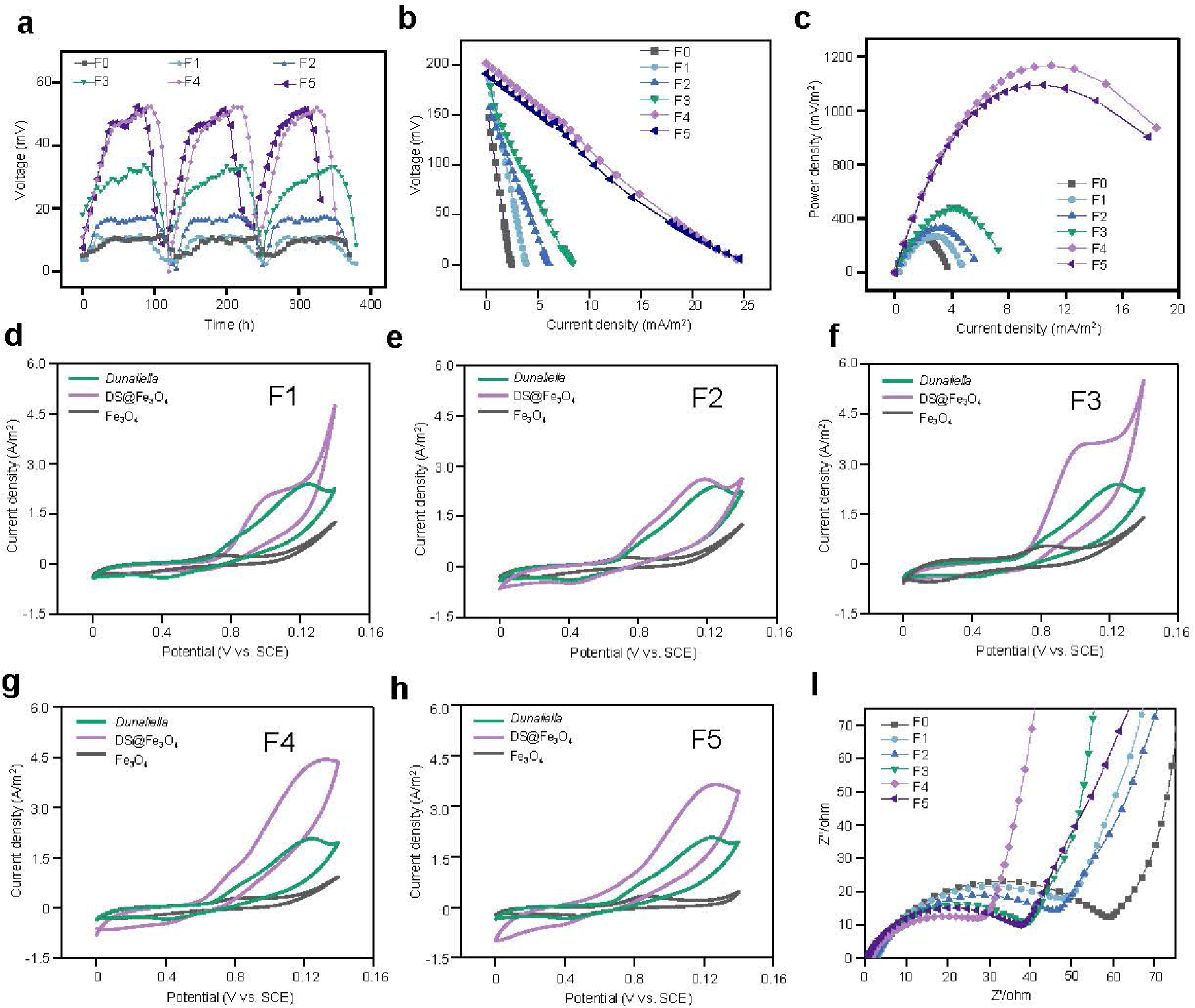
Electrochemical analysis of *Dunaliella* BPVs with different Fe_3_O_4_ coating concentrations. **a.** *Dunaliella* BPVs with different Fe_3_O_4_-coated concentrations versus time curves for long-term stability and repeated cycling tests (F0: 0 mg/mL, F1: 0.25 mg/mL, F2: 0.5 mg/mL, F3: 1.0 mg/mL, F4: 2 mg/mL, F5: 4 mg/mL). Polarization curves (**b**) and power output curves (**c**) of *Dunaliella* BPVs with different Fe_3_O_4_-coated concentrations. **d-h**. CV curves of *Dunaliella* BPVs with different Fe_3_O_4_-coated concentrations. **d**, F1 (0.25 mg/mL); **e**, F2 (0.5 mg/mL); **f**, F3 (1 mg/mL); **g**, F4 (2 mg/mL); **h**, F5 (4 mg/mL). CV curves at a scan rate of 10 mV·s^−1^. **i**. EIS spectra of *Dunaliella* BPVs with different Fe_3_O_4_-coated concentrations. The *Dunaliella* cells were set to a final working OD_630_ = 0.7.

The power density and polarization curves for *Dunaliella* BPVs coated with varying concentrations of Fe_3_O_4_ were shown **Fig. 3b**. The polarization curves demonstrated that the maximum OCV for F0, F1, F2, F3, F4 and F5 BPVs were 198.29, 190.15, 185.14, 183.23, 194.87, and 200.51 mV, respectively. These results indicated that the OCV were similar across different Fe_3_O_4_ concentrations, showing that *Dunaliella* maintained a comparable electricity generation capacity at the same biomass level (**Supplementary Table S5, 6**). As the shell concentration increased, the slope of the fitted curves decreased, suggesting a reduction in the apparent internal resistance and an enhancement in electron transfer efficiency of the *Dunaliella* BPVs. The power density curves revealed that the maximum power densities for F0, F1, F2, F3, F4 and F5 BPVs built were 190.27, 201.10, 245.48, 363.15, 1150.64, and 1041.77 mW/m^2^, respectively. This showed that the performance of *Dunaliella* BPVs improved with increasing Fe_3_O_4_ concentration, reaching its peak at 2 mg/mL. At this concentration, F4 produced a maximum power density of 1150.64 mW/m^2^, which was 5.79 times higher than that of a native *Dunaliella* BPV (190.27 ± 1.66 mW/m^2^). However, as the Fe_3_O_4_ concentration continued to rise, the power density began to decrease, dropping to 1067.30 mW/m^2^ at 4 mg/mL, which was still 6.04 times that of the native *Dunaliella* BPV. This indicated that the BPV with 2 mg/mL Fe_3_O_4_-coated *Dunaliella* exhibited superior electrochemical performance.

### Electrochemical performance of *Dunaliella* BPVs with different Fe_3_O_4_-coated concentrations

To assess the impact of Fe_3_O_4_ nanoparticles on the electron transfer efficiency of *Dunaliella*, CV analyses were performed on bioanode electrodes of BPVs containing different concentrations of Fe_3_O_4_-coated *Dunaliella*. Under varying concentrations of Fe_3_O_4_, the electrochemical activity of the *Dunaliella* bioanodes changed significantly. At lower Fe_3_O_4_ concentrations of 0.25 and 0.5 mg/mL, the oxidation peak currents in the CV curves were similar to those without Fe_3_O_4_, indicating a minimal impact on the anode’s electrochemical activity. However, at a concentration of 1 mg/mL, both the size of the oxidation peak and the peak current increased substantially, signifying a marked enhancement in the bioanode’s electrochemical activity under the influence of 1 mg/mL Fe_3_O_4_. The strongest electrochemical activity of the shell-coated *Dunaliella* bioanodes corresponded to a Fe_3_O_4_ concentration of 2 mg/mL. Increasing the Fe_3_O_4_ concentration beyond 2 mg/mL led to a decrease in electrochemical activity, suggesting that higher concentrations of Fe_3_O_4_ might adversely affect the microbial growth and metabolism, thereby reducing the electron transfer capability on the bioanode surface.

To evaluate changes in the bioanode conductivity of *Dunaliella* cells before and after coating with varying concentrations of Fe_3_O_4_, EIS was used to measure the conductivity of the biofilm. Nyquist plots for BPV electrodes with different concentrations of DS@Fe_3_O_4_ were shown in **Fig3. i**, with the fitting circuit data provided in **Supplementary Table S7**. The native *Dunaliella* BPV exhibited the highest charge transfer resistance, whereas the DS@Fe_3_O_4_ bioanodes showed significantly reduced impedance, with the sequence of resistance values being F0 > F1 > F2 > F3 > F5 > F4. After coating the *Dunaliella* cells with different concentrations of Fe_3_O_4_, it was observed that *R*_ct_ gradually decreased as the coating concentration increased from 0 to 1 mg/mL, dropping from 61.8 to 41.03 Ω. The lowest *R*_ct_, at 22.72 Ω, occurred at a Fe_3_O_4_ concentration of 2 mg/mL, making the native *Dunaliella* bioanode 2.72 times higher. However, as the Fe_3_O_4_ concentration increased further, *R*_ct_ began to rise again, reaching 36.44 Ω at 4 mg/mL, which was 1.69 times that of the *Dunaliella* bioanode.

In summary, Fe_3_O_4_ concentration of 2 mg/mL provided the highest electron transfer rate for the shell-coated *Dunaliella* BPV. This demonstrated the potential of Fe_3_O_4_ coating to enhance conductivity on the surface of algal bio-units, although there is an optimal coating load. These findings can promote the application of these bio-units in electron-related fields.

### SiO_2_ shell-coating reduces the electron transfer efficiency of Fe_3_O_4_ in *Dunaliella* BPVs

To further explore the mechanism of Fe_3_O_4_ in *Dunaliella* BPVs, we performed shell-coating of Fe_3_O_4_with a layer of SiO_2_ (Fe_3_O_4_@SiO_2_, **Supplementary Fig. S8**), and then applied Fe_3_O_4_@SiO_2_ for coating *Dunaliella*. In **Fig. 4a**, compared to DS@Fe_3_O_4_, the DS@Fe_3_O_4_@SiO_2_ solution remained green, and the cells did not exhibit significant sedimentation. Under the microscope, Fe_3_O_4_@SiO_2_ did not adhere well to the *Dunaliella* cells, with many particles dispersed around the cells.

**Figure 4.**
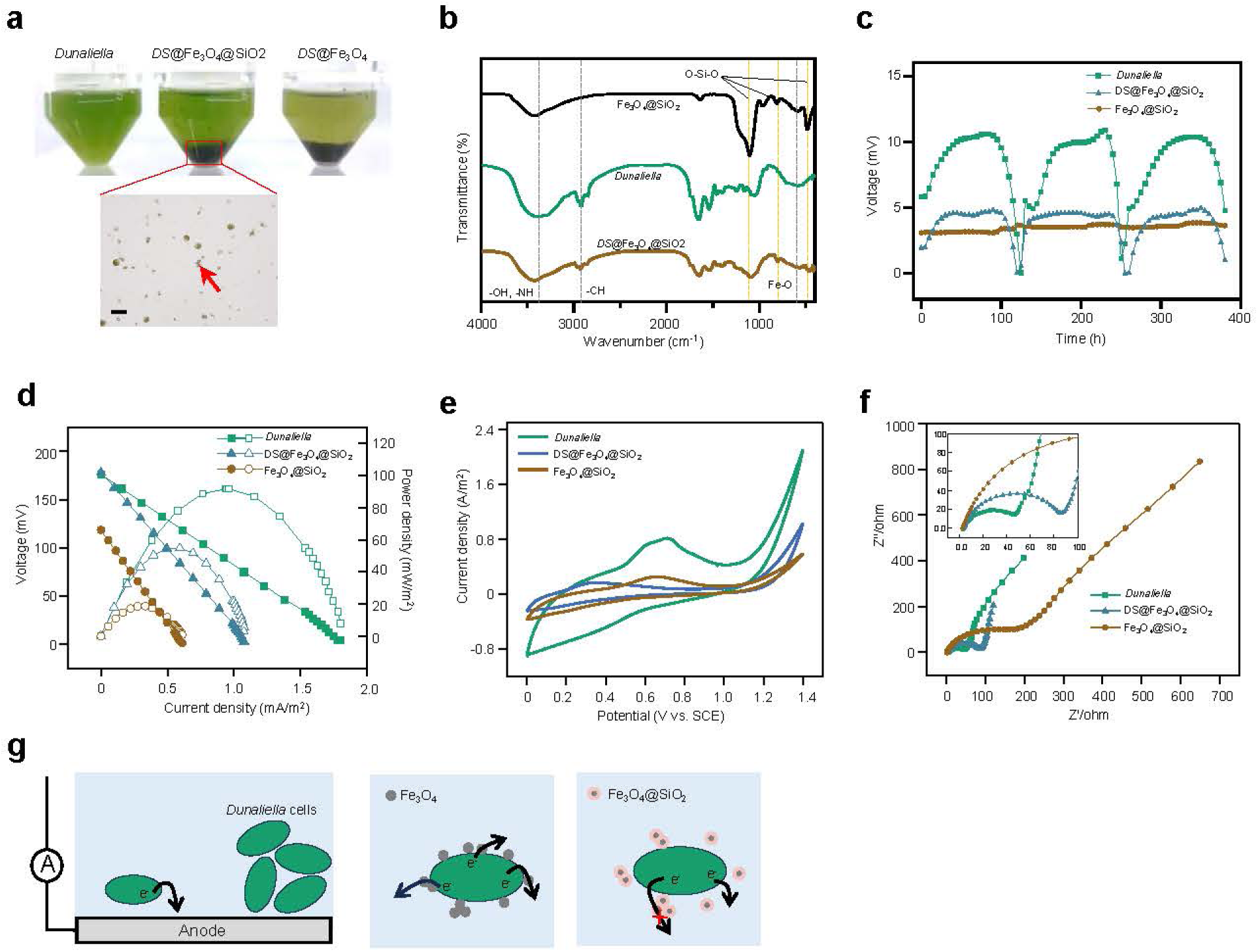
Characterization and performance analysis of Fe_3_O_4_@SiO_2_-coated *Dunaliella* BPVs. **a.** Algal fluid and optical microscope of *Dunaliella* cells coated with Fe_3_O_4_@SiO_2_. **b.** Infrared spectra of Fe_3_O_4_@SiO_2_, native and Fe_3_O_4_-coated *Dunaliella* cells. **c.** Native *Dunaliella*, Fe_3_O_4_@SiO_2_, and DS@Fe_3_O_4_@SiO_2_ BPVs versus time curves for long-term stability and repeated cycling tests. Polarization curves and power output curves (**d**), CV curves (**e**), and EIS spectra (**f**), of native *Dunaliella,* Fe_3_O_4_@SiO_2_, and DS@Fe_3_O_4_@SiO_2_ BPVs. **g.** Schematic diagram of electron transfer in native, Fe_3_O_4_-coated, and Fe_3_O_4_@SiO_2_-coated *Dunaliella* BPVs. The Fe_3_O_4_ coating enhances the efficiency of electrons to transfer to the anode of the BPV. The addition of SiO_2_ blocks the transfer of electrons through Fe_3_O_4_ to the anode.

To explore the impact of Fe_3_O_4_@SiO_2_ coating on the electrical output of *Dunaliella* BPVs used as power-generating anodes, the output voltage of DS@Fe_3_O_4_@SiO_2_ during stable operation over three cycles was measured, as illustrated in **Fig. 4b**. The bioanode of native *Dunaliella* BPV exhibited higher output voltage than those modified with Fe_3_O_4_@SiO_2_. The maximum output voltage for DS@Fe_3_O_4_@SiO_2_ was 4.80 ± 0.10 mV, which is a 54.72% decrease compared to the native *Dunaliella* BPV output voltage of 10.60 ± 0.15 mV. Both the DS@Fe_3_O_4_@SiO_2_ and native *Dunaliella* BPVs exhibited similar OCV (**Fig. 4c** and **Supplementary Table S8, 9**), suggesting comparable power-generating capabilities. However, the apparent internal resistance was significantly higher for the DS@Fe_3_O_4_@SiO_2_ BPV at 161.31 Ω compared to 94.21 Ω for the native *Dunaliella* BPV, suggesting that the SiO_2_ layer substantially increases material resistance, thereby impeding electron transfer. The maximum power density generated by the *Dunaliella* BPV coated with Fe_3_O_4_@SiO_2_ was 55.89±0.48 mW/m^2^, which is a 38.12% decrease from the 90.32±0.24 mW/m^2^ observed with the native *Dunaliella* anode. Interestingly, the addition of SiO_2_ to the Fe_3_O_4_ shell seems to block electron transfer from the surface of the algae to the electrode.

To further investigate the performance of DS@Fe3O4@SiO2 BPV, CV was utilized (**Fig. 4d**). The bioanode of the uncoated native *Dunaliella* exhibited an oxidation peak at approximately 0.7 V (vs. SCE). In contrast, the DS@Fe_3_O_4_@SiO_2_ bioanode showed a weaker oxidation peak current. EIS measurements and equivalent circuit modeling provided quantitative data to assess the impact of Fe_3_O_4_@SiO_2_ on the performance of *Dunaliella* BPVs (**Fig. 4e** and **Supplementary Table S10**). The sequence of electrode impedance was as follows: Fe_3_O_4_@SiO_2_ (256.1 Ω) > DS@ Fe_3_O_4_@SiO_2_ (91.66 Ω) > *Dunaliella* (45.28 Ω), indicating that the Fe_3_O_4_@SiO_2_ coating increases charge transfer resistance compared to native *Dunaliella* BPVs. This indicated that the slope was much steeper for BPVs using DS@Fe_3_O_4_@SiO_2_, correlating with higher internal resistance and consistent with the polarization results.

In conclusion, Fe_3_O_4_ serves as a single-cell electron collector enhancing the non-biological/biological interface electron transfer efficiency and BES performance, potentially breaking through the limitations of electron transfer at these interfaces (**Fig. 4f**). Enhanced current output can be attributed to improved electron transport at the cell/electrode interface and through the cellular network. The interconnection of nanoparticles with *Dunaliella* cells represents a unique and promising direction in BPV research, potentially enhancing our understanding of electron transfer processes in these biological systems and overcoming a key limitation in BPVs by constructing a connected 3D cellular network.

## Discussion

This study introduced a novel method to enhance *Dunaliella*-based BPVs by coating the algae with Fe_3_O_4_ and Al_2_O_3_ nanoparticles. The Fe_3_O_4_ coating significantly improved electron transfer efficiency and overall BPV performance, as evidenced by higher peak voltages and power densities compared to native *Dunaliella* BPVs. This enhancement is attributed to the excellent conductivity and biocompatibility of Fe_3_O_4_, which facilitated efficient electron transfer from the cells to the electrode surface.

Optimal Fe_3_O_4_ concentration for coating was found to be 2 mg/mL, yielding the highest performance. However, higher concentrations decreased performance, likely due to cytotoxic effects or mechanical stress on the cells. In contrast, DS@Al_2_O_3_ BPVs showed only marginal improvements, suggesting limited conductivity, potential cytotoxicity, and the inherent non-conductive nature of Al_2_O_3_.

Incorporating a SiO_2_ layer on Fe_3_O_4_ impeded electron transfer, resulting in lower voltages and power densities due to increased internal resistance and reduced Fe_3_O_4_’s attachment to *Dunaliella* cells. Electrochemical analyses confirmed these findings, showing enhanced electrochemical activity and reduced charge transfer resistance for Fe_3_O_4_-coated *Dunaliella*, but decreased performance when further coated with SiO_2_.

In summary, Fe_3_O_4_ coatings substantially improve the electron transfer efficiency and performance of *Dunaliella*-based BPVs. This approach holds promise for advancing bioelectrochemical systems for energy and environmental applications. Future research should focus on optimizing Fe_3_O_4_ concentrations, exploring alternative coatings, and evaluating long-term stability and scalability for practical implementation.

## Methods

### Culture conditions

The strain *Dunaliella salina* CCAP 19/18 was procured from the Culture Collection of Algae and Protozoa (CCAP), Scotland, United Kingdom. It was cultivated in a 2ASW (artificial seawater) medium, with a saline concentration of 1.1 M NaCl as detailed in **Supplementary table S11**, **S12**^50,51^. The algal cultures were maintained at 26°C within a growth chamber, under a luminous flux of 8, 000 cd·sr/m^2^ supplied by cool-white fluorescent lamps, following a 16/8 h light/dark cycle. The optical density (OD) of the algal cultures was ascertained using an Agilent spectrophotometer (USA) at a wavelength of 630 nm (OD_630_). Cell populations within the algal cultures were quantified employing a Zeiss Axioplan microscope (Germany), equipped with a 40× (N.A. 0.75) plan-apochromatic objective and a 100-W mercury lamp, along with a hemocytometer for precise counting. See **Supplementary Note 3** for full details.

### Single-Cell nanoencapsulation

The procedure commences by centrifuging the sample at 1,000 rpm for 3 minutes to collect *D. salina* cells, followed by two washes using a standard algal culture medium (1.1 M NaCl). Subsequently, the cells are resuspended in a dispersal algal culture medium (pH = 7.0), with the cell concentration adjusted to 10^8^ cells/mL (by centrifuging 9 mL of algal suspension at 1,000 rpm for 3 minutes, discarding the supernatant, and resuspending the cells in 3 mL of culture medium). Nano-sized Al_2_O_3_ or Fe_3_O_4_ was then introduced to achieve a final concentration of 0, 0.25, 0.5, 1, 2, and 4 mg/mL (prepared using algal culture medium and pre-dispersed with ultrasonication), and the mixture was placed in a shaking incubator at 120 rpm for 3 hours. Lastly, a portion of the cells is allocated for further culturing, while another portion is prepared for electron microscopy examination.

As for Ds@Fe_3_O_4_@SiO_2_, Nanoparticles of Fe_3_O_4_@SiO_2_, synthesized chemically (**Supplementary note 4**), were added to a dispersion of *Dunaliella* cells to achieve a concentration of 0.25 mg/mL. After shaking at 120 rpm for 30 minutes on a shaker, the mixture was ready for subsequent cultivation.

### Core-shell Characterizations

Using optical microscopy (Nikon Eclipse 50i, Japan), scanning electron microscopy (SEM), and transmission electron microscopy (TEM), the morphology and structure of *Dunaliella* samples before and after shell-coating were characterized. The specific procedures were as follows: the coated *Dunaliella* samples were first fixed in 2.5% glutaraldehyde solution for electron microscopy at 4°C for 24 hours. After fixation, the cells were washed three times with ultrapure water to remove the fixative.

SEM observations and energy-dispersive X-ray spectroscopy (EDX) analyses were performed using a Merlin (Zeiss, Germany) and X-MaxN20 dual detector system. Silicon substrates were used for SEM imaging, and aluminum substrates were used for EDX analysis. The samples were dropped onto silicon and aluminum substrates, respectively, and left to air dry at 30°C for 24 hours. Before testing, the samples were sputter-coated with gold.

TEM observations were performed using FEI Talos F200x (USA). The samples fixed in 2.5% glutaraldehyde were treated with OsO_4_ and K_2_Cr_2_O_7_, dehydrated with acetone/ethanol, embedded in resin, and then lifted onto copper grids (with ultrathin carbon film) for observation under the transmission electron microscope.

Fourier Transform Infrared Spectroscopy (FTIR) analysis of *Dunaliella* powder samples before and after shell-coating, along with nanoparticle powders at the same concentrations, was performed using a Thermo Scientific iN10 spectrometer (USA).

Native *Dunaliella*, shell-coated *Dunaliella*, and their nanoparticle suspensions at the same cell concentration (1×10^6^ cells/mL) were dropped onto acrylic plates and allowed to dry naturally. The ultraviolet-visible transmission spectra of the *Dunaliella* samples before and after shell-coating, as well as of the nanoparticle suspensions at the same concentration, were measured using a PE Lambda 750 UV-Vis spectrophotometer (USA).

### Chlorophyll fluorescence analysis

To ensure that the PSII reaction centers were open, every 12 hours, 3 mL of native *Dunaliella* cells and Fe_3_O_4_-coated *Dunaliella* cells (DS@Fe_3_O_4_) were placed in sample tubes and kept in the dark for 20 minutes, thoroughly oxidizing the photosynthetic electron transport chain. The steady-state yields of chlorophyll fluorescence were measured with an AquaPen Fluorometer (AP-110, Czech Republic) set to ’Fluo’ mode. The minimal fluorescence at open PSII centers in the dark-adapted state (*Fo*) was excited by a weak measuring light (650 nm). A saturating pulse of red light (2, 400 μmol·m^-2^·s^-1^) was applied to determine the maximal fluorescence at closed PSII centers in the dark-adapted state (*Fm*). The maximal quantum yield of PSII (*Fv/Fm*) was calculated as (*Fm - Fo*) / *Fm*^52^.

### Bioelectronic device setup

The BPV configuration was a dual-chamber made of plexiglass, with both the anode and cathode chambers having a volume of 100 mL each, separated by a Nafion 177 proton exchange membrane (DuPont, USA). The pretreated carbon cloth (CeTech GF020, **Supplementary note 5**) was placed in the two chambers, serving as both anode and cathode. The electricity-generating *Dunaliella* suspension was introduced into the anode chamber, and the cathode electron acceptor electrolyte was introduced into the cathode chamber. The cathode and anode carbon cloth electrodes were connected externally with titanium wire (1 mm dia.) and were linked to a decade box ZX21 (Shanghai Domore, China) via conducting wires to form a closed circuit (Setting the resistance to 1 kΩ). The digital multimeter data logger VC86E (Victor, China) was connected at both ends of the circuit to record data with VICTOR 86E V4.0 software. The real-time output voltage of the BPV reactor was automatically recorded, with voltage values logged every 60 minutes. The BPV reactor was placed in an illuminated incubator (HYG-C, TCSYSB, China), with the temperature controlled at 23 ± 1°C, and the light intensity set to 8, 000 cd·sr/m^2^.

### Comparison of BPV performance with different cathode electron acceptor electrolyte

The experiment selected K_3_[Fe(CN)_6_], KMnO_4_, and K_2_S_2_O_8_ as cathode electron acceptors. The anolyte consisted of *Dunaliella* medium, inoculated with *Dunaliella* cells to achieve a final working OD_630_ = 1.2. The catholyte consisted of potassium persulfate solution (K_2_S_2_O_8_, 50 mmol/L), potassium permanganate solution (KMnO_4_, 50 mmol/L), and potassium ferricyanide solution (K_3_[Fe(CN)_6_], 50 mmol/L). The electrochemical reactions were shown in the following equations.

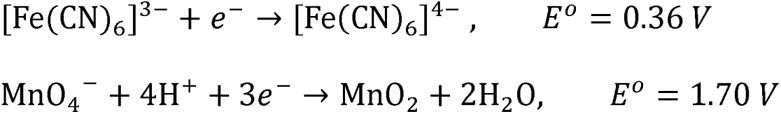

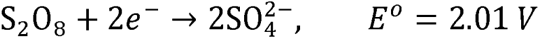

### Comparison of BPV performance with Fe_3_O_4_ and Al_2_O_3_ shell-coated *Dunaliella*

K_2_S_2_O_8_ served as the cathode electron acceptor in the construction of a shell-coated *Dunaliella* BPV. The anolyte consisted of *Dunaliella* medium inoculated with shell-coated (nano Fe_3_O_4_ or Al_2_O_3_) or uncoated *Dunaliella* cells to achieve a final working OD_630_ = 1.0. Due to the shift in the UV absorption peak of the shell-coated algae cells, to maintain a consistent number of algae cells, the OD_630_ was strictly controlled before preparation. It was important to note that OD_630_ measures uncoated algae cells.

### Comparison of BPV performance with ladder concentration Fe_3_O_4_ shell-coated *Dunaliella*

*Dunaliella* cells (OD_630_ = 0.7) were coated on the cell surface with Fe_3_O_4_ nanoparticles at concentrations of 0, 0.25, 0.5, 1, 2, and 4 mg/mL. BPVs were constructed using *Dunaliella* cells coated with different concentrations of Fe_3_O_4_ nanoparticles, designated as F0 (0 mg/mL), F1 (0.25 mg/mL), F2 (0.5 mg/mL), F3 (1 mg/mL), F4 (2 mg/mL), and F5 (4 mg/mL). The anolyte consisted of *Dunaliella* cells and *Dunaliella* cells coated with different concentrations of Fe_3_O_4_, while the catholyte was 50 mmol/L K_2_S_2_O_8_ solution.

### Electrochemical surface area (ECSA) measurement

Cyclic voltammetry (CV) is an electrochemical evaluation method, effectively used to assess the performance of the electrochemical surface area (ECSA) in batteries. In this experiment, CV tests were conducted using a constructed three-electrode system at an electrochemical workstation CS310 (CORRTEST, China). The working electrode in the three-electrode system consisted of a biofilm anode formed by *Dunaliella* cells and their nano-material-coated counterparts attached to carbon cloth (CC), with a counter electrode as the cathode, and a saturated calomel electrode as the reference electrode CHI150 (+0.2412V vs. SHE) (Chenhua, China) monitoring the potentials of the anode and cathode. During the cyclic voltammetry test, CV scans the voltage back and forth from a lower value to a higher value, recording the current response and displaying it on the curve of the voltammogram.

### Electrochemical impedance spectroscopy (EIS) measurement

Electrochemical impedance spectroscopy (EIS) is another electrochemical test used to evaluate the electrochemical performance of electrodes. The constructed three-electrode system was tested using the impedance mode function at an electrochemical workstation CS310 (CORRTEST, China). In this setup, the working electrode was the biofilm anode made of *Dunaliella* or its coated cells, the counter electrode was carbon cloth, and the reference electrode was a saturated calomel electrode CHI150 (Chenhua, China). The AC impedance frequency parameters were set from 100 kHz to 10 mHz with an amplitude of 5 mV. The EIS test results were imported into Zview software and fitted using a Randles’ equivalent circuit diagram, with the components of the circuit diagram consistent with the BPV circuit model. The fitted data obtained were used to analyze the results from the electrochemical system tests and the physicochemical properties of the BPV electrodes. For the full BPV, the EIS test was conducted after the BPV output voltage had stabilized at a high plateau.

## Author contributions

HC, conceptualization and experiment design, visualization, wrote the original draft; JW, conducted some experiments, data analysis; JJ, manuscript revision and polish, project administration, funding acquisition. All authors reviewed and provided input on the manuscript.

## Acknowledgement

This project was supported by the National Natural Science Foundation of China (32372286 and 32072201), Guangdong Basic and Applied Basic Research Foundation (2023A1515011967 and 2023A1515012223).

## Supplementary information

In separated documents including supplementary information and tables.

## Data availability

Data are available from the corresponding author upon reasonable request.

