## Supplementary Note for "Fe_3_O_4_ nanoparticles shell amplify charge-extraction efficiency in *Dunaliella* photovoltaics"

### Supplementary note 1, Zeta potential

**Zeta potential measurement**

The Zetasizer Nano S (Malvern, UK) was used to measure the zeta potentials of native *Dunaliella* cells, as well as *Dunaliella* cells coated with $Fe_3O_4$, $Al_2O_3$ and $Fe_3O_4@SiO_2$. During testing, the cell densities for both the native and coated *Dunaliella* cells were approximately $2\times10^6$ cells/mL, suspended in a *Dunaliella* medium with a pH of 7.0.

**Zeta potential analysis**

Changes in the surface Zeta potential of *Dunaliella* before and after shell-coating can be analyzed to explore the mechanisms of cell shell-coating (**Supplementary Fig. S2**). The Zeta potential of the original *Dunaliella* before mineralization was -31.07 ± 1.25 mV, and for $Fe_3O_4$ nanoparticles, it was 1.35 ± 0.13 mV. After dispersing the cells into the nanoparticles to coat the cell surface, the system's Zeta potential significantly increased to -25.63 ± 1.54 mV. Nanoparticles with opposite charges tend to neutralize each other's charge[3]. According to previous infrared analyses, groups like -OH, C=O, and N-H on the surface of *Dunaliella* cells cause them to carry a negative charge in water or culture media due to dehydrogenation through protonation[4]. This indicates that interactions occurred between the $Fe_3O_4$ nanoparticles and the cells, and the magnetic nanoparticles were primarily adsorbed onto the cell surface through electrostatic attraction.

#### Supplementary note 2, Formation of DS@$Al_2O_3$ core-shell structure

To compare with the highly conductive $Fe_3O_4$, we selected $Al_2O_3$ nanoparticles, which are poorly conductive, and investigated their interaction with *Dunaliella* cells through direct introduction. We discovered that $Al_2O_3$ nanoparticles also have self-assembly properties (**Supplementary Fig. S6**). Typically, under neutral pH conditions of their culture environment, *Dunaliella* cells exhibited a negatively charged surface, while $Al_2O_3$ nanoparticles carried a positive charge (**Supplementary Fig. S2**). SEM observations showed that the surface of native *Dunaliella* cells is smooth, whereas in the $Al_2O_3$ system, DS@$Al_2O_3$ had numerous particles deposited on the surface, and $Al_2O_3$ completely enveloped the *Dunaliella*, forming a rough algal shell.

TEM cross-sectional images revealed that $Al_2O_3$ nanoparticles were only coated on the cell membrane surface and did not penetrate inside the cells. The structural integrity of the *Dunaliella* cells was maintained, likely because the particle size of the $Al_2O_3$ nanoparticles was much larger than the pore size of the cell membrane surface. $Al_2O_3$ particles tightly adhered to the outer layer of the *Dunaliella* cells.

#### Supplementary note 3, *Dunaliella* cell numbers measurement

Due to the rapid aggregation phenomenon of *Dunaliella* cells coated with $Fe_3O_4$ and $Al_2O_3$ nanoparticles, it becomes difficult to measure the number of *Dunaliella* cells using UV spectrophotometry. Therefore, we used optical microscopy and random counting to statistically determine the number of *Dunaliella* cells. We took 10 μL of well-mixed samples with different concentrations (0, 0.25 mg/mL, 0.5 mg/mL, 1 mg/mL, 2 mg/mL, and 4 mg/mL) of $Fe_3O_4$ and $Al_2O_3$ and placed them on a coverslip. A small amount of 2.5% glutaraldehyde fixative was added, covered with a coverslip, and observed under an optical microscope. For each sample, three replicates were prepared, and five fields were randomly selected for cell counting.

#### Supplementary note 4, Synthesis of nanoparticles

**$Fe_3O_4$**

Initially, 0.99 g of $FeCl_2 \cdot 4H_2O$ and 2.7 g of $FeCl_3 \cdot 6H_2O$ were dissolved in a four-neck flask containing 100 mL of distilled water. This mixture was stirred vigorously under a nitrogen atmosphere at 80°C. Subsequently, 10 mL of $NH_4OH$ (25 wt.%) was introduced into the solution. Continuous stirring was maintained for an additional 30 minutes, during which the solution transitioned in color from light brown to black, indicating the formation of magnetite. The resultant magnetite nanoparticles were separated from the mixture using a small permanent magnet, and were rinsed four times with distilled water to ensure purity.

**$Fe_3O_4$@$SiO_2$**

An improved Stöber chemical synthesis method enabled the deposition of a $SiO_2$ layer on the surface of $Fe_3O_4$, preparing $Fe_3O_4$@$SiO_2$ particles with a core-shell structure[1]. A mixture of ethanol (160 mL) and ammonia solution (16 mL) was added to deionized water (24 mL) and mixed, then $Fe_3O_4$ particles (260 mg) were added. The mixture was transferred to a flask and vigorously mechanically stirred at 50°C for 10 minutes to form a uniformly distributed system. Subsequently, a total of 400 μL of TEOS (tetraethoxysilane, Sigma) was added to the mixture at a rate of 200 μL/30 min. After 30 minutes of reaction, the $Fe_3O_4$@$SiO_2$ particles were separated with a magnet. The product was washed with deionized water and ethanol. The mixture was dried in a vacuum oven at 60°C to obtain the $Fe_3O_4$@$SiO_2$ material.

The morphology of iron oxide and $Fe_3O_4$@$SiO_2$ nanoparticle materials was characterized with a FEI Talos F200x transmission electron microscope (USA). Fourier Transform Infrared Spectroscopy (FTIR) analysis of the $Fe_3O_4$ and $Fe_3O_4$@$SiO_2$

powder samples was conducted with a Thermo Scientific iN10 FTIR spectrometer (USA).

**Transmission electron microscopy characterization of $Fe_3O_4$ and $Fe_3O_4$@$SiO_2$ nanoparticles**

To further analyze the structure and size of the synthesized $Fe_3O_4$ and its core-shell $Fe_3O_4$@$SiO_2$, we characterized the materials using a transmission electron microscope (TEM). **Supplementary Fig. S8a** showed spherical magnetic nanoparticles of $Fe_3O_4$ with smooth surfaces and a particle diameter of approximately 10-20 nm. **Supplementary Fig. S8b** displayed the $Fe_3O_4$@$SiO_2$, where a black $Fe_3O_4$ nanoparticle core and a gray silicon dioxide shell of significant thickness were visible, forming a core-shell-like structure with a diameter of about 20 nm. Each nanosilica sphere encapsulates one or more magnetite nanocrystals, with the silicon dioxide shell thickness around 5-10 nm. However, aggregation/clustering of individual $Fe_3O_4$@$SiO_2$ was observed, and aggregation may occur during TEM sample preparation and under other observed drying conditions[2].

#### Supplementary note 5, BPVs material pretreatment

##### Pretreatment of the proton exchange membrane

The Nafion 117 membrane (DuPont, USA) was pretreated before use to achieve better electron transfer. The membrane was placed in a 30% hydrogen peroxide solution and boiled in a water bath at 80°C for 1 hour. It was then soaked in deionized water for 0.5 hours. The membrane was boiled in 5% dilute sulfuric acid at 100°C for 1 hour, followed by boiling in deionized water at 60°C for 0.5 hours. Finally, it was soaked in deionized water for another 0.5 hours. After the Nafion 117 membrane was removed, it was dried and stored.

##### Pretreatment of carbon cloth

The carbon cloth was cut into circles with a diameter of 5.0 cm, then soaked and sonicated in 1 mol/L HCl for 1 hour, followed by soaking and sonication in deionized water for 1 hour. It was then soaked in acetone for 1 hour before being soaked in deionized water for 1 hour to remove surface impurities, and finally, it was dried for later use.

#### Supplementary note 6, Cyclic voltammetry (CV) scans

By performing cyclic voltammetry (CV) scans at different scan rates on *Dunaliella* and its shell-coated BPVs, the anode CV curves measured in the native *Dunaliella* BPV, DS@$Fe_3O_4$ BPV, and DS@$Al_2O_3$ BPV varied with scan rates (10, 8.0, 6.0, 4.0, 2.0 mV/s), as shown in **Supplementary Fig. S9a-c.**

Theoretically, if the absolute value of the peak current intensity (Ip) was linearly related to the scan rate (*v*) (**Supplementary Fig. S9d**), it could be initially judged that the power generation mechanism was an adsorption process, where electricity was generated through the formation of a thick biofilm. If Ip was linearly related to the square root of *v*, it could be judged that the power generation process was a diffusion process, where electricity was indirectly generated through the formation of electron carriers. The Ip of *Dunaliella* MFC at different scan rates showed a good linear correlation with *v*, indicating that its power generation mechanism was an adsorption process. Therefore, it could be initially judged that *Dunaliella* transferred extracellular electrons through direct contact between the cells and the electrode. Related studies on green microalgae *Chlorella* MFCs mentioned that the algae responsible for biocatalysis in the cell were not the suspended cells in the algal solution but the microalgae biofilm formed on the electrode.

As shown in **Supplementary Fig. S9d**, the peak currents of both systems were approximately proportional to the scan rate, with the slope of the DS@$Fe_3O_4$ BPV (1.05) being about 4.56 times that of the native *Dunaliella* BPV (0.23). The slope of the DS@$Al_2O_3$ BPV (0.26), which had poor conductivity, was similar to that of the native *Dunaliella* BPV. Under the same scan rate conditions, the response current of the DS@$Fe_3O_4$ BPV bio-anode was much greater than that of the control group with the native *Dunaliella* bio-anode, indicating that the DS@$Fe_3O_4$ BPV could accelerate electron transfer from the cell to the electrode surface, thus accelerating the catalytic oxygen reduction reaction at the cathode. Therefore, the anode reaction in this study was mainly controlled by adsorption, and using $Fe_3O_4$ to shell-coat *Dunaliella* cells significantly enhanced the adsorption capacity and electron transfer efficiency of the anode cells.

### Supplementary Figures

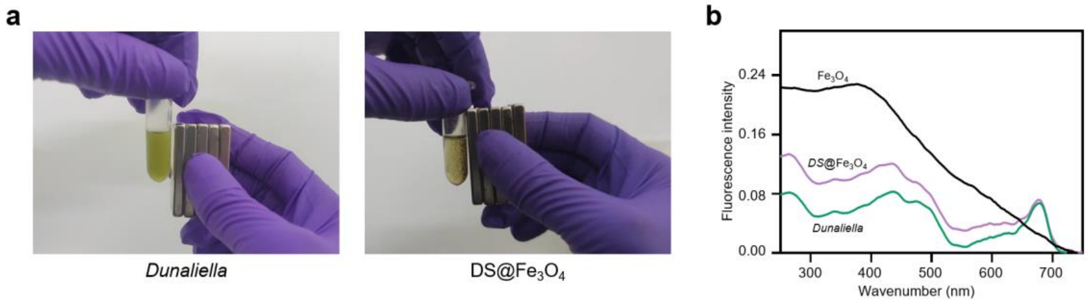


**Supplementary Fig. S1. a**. The magnetic properties of DS@$Fe_3O_4$ cells. DS@$Fe_3O_4$ cells aggregate to the right under the attraction of a magnet. The magnetic properties of DS@$Fe_3O_4$ cells were also demonstrated in the **Video**. **b**. UV-vis absorption spectrum of *Dunaliella* cell before and after $Al_2O_3$ coating.

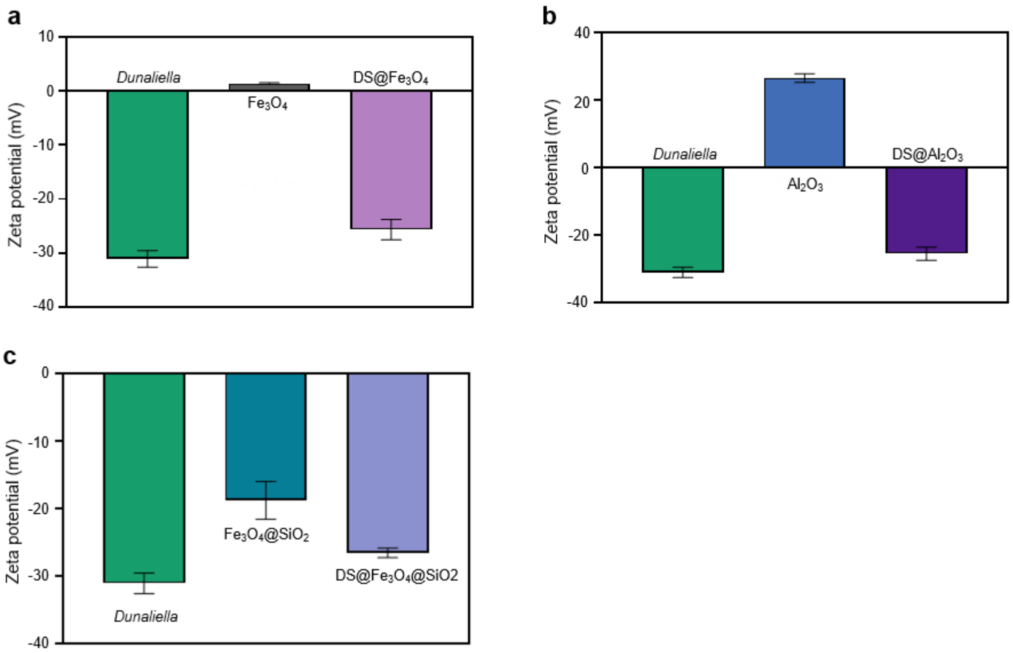


**Supplementary Fig. S2.** The Zeta potential of *Dunaliella* before and after coating with $Fe_3O_4$ (**a**), $Al_2O$ (**b**)$_3$, $Fe_3O_4$@$SiO_2$ (**c**) nanoparticles and the Zeta potential of the nanoparticles themselves. pH=7.0. N = 3 biologically independent samples and data are presented as mean ± s.d..

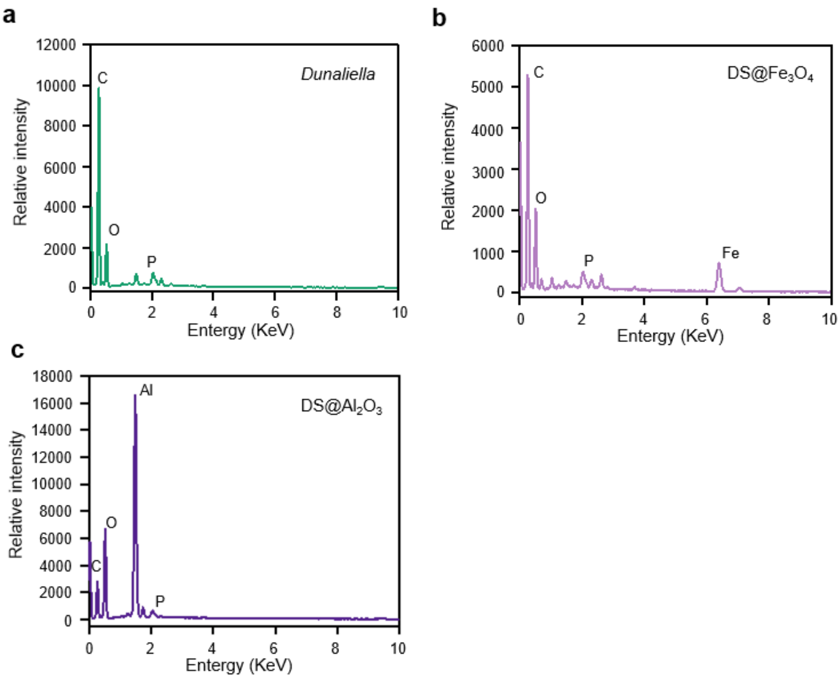


**Supplementary Fig. S3. EDX profile of native *Dunaliella* (a), DS@$Fe_3O_4$ (b) and DS@$Al_2O_3$ (c) cells.** Using Energy-dispersive X-ray spectroscopy (EDX) for further characterization of native *Dunaliella* cells, DS@$Fe_3O_4$, and DS@$Al_2O_3$ cells, it was found that they all had high C element content. However, only DS@$Fe_3O_4$ cells contained Fe and only DS@$Al_2O_3$ cells contained Al, indicating that $Fe_3O_4$ and $Al_2O_3$ were successfully coated onto the surface of *Dunaliella* cells, and the core-shell structures of DS@$Fe_3O_4$ and DS@$Al_2O_3$ were successfully constructed.

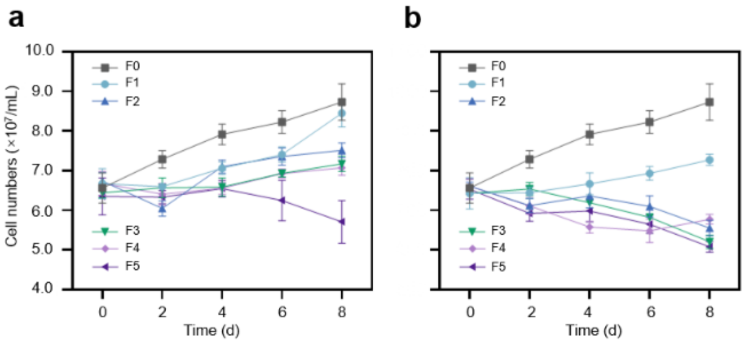


**Supplementary Fig. S4.** Cell growth of *Dunaliella* cells coated with $Fe_3O_4$ (**a**) and $Al_2O_3$ (**b**) at different concentrations (F0: 0 mg/mL; F1: 0.25 mg/mL; F2: 0.5 mg/mL; F3: 1.0 mg/mL; F4: 2 mg/mL; F5: 4 mg/mL). The growth curves of native *Dunaliella*, DS@$Fe_3O_4$, and DS@$Al_2O_3$ cells indicated that high concentrations (4 mg/mL) of $Fe_3O_4$ and $Al_2O_3$ suppressed cell division. Additionally, $Al_2O_3$ concentrations ranging from 0.5 to 0.4 mg/mL also inhibited *Dunaliella* growth.

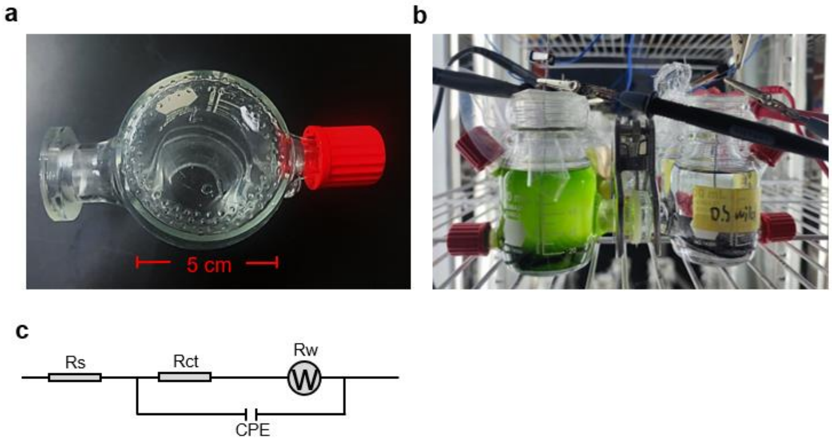


**Supplementary Fig. S5. The setup of the dual-chamber *Dunaliella* biophotovoltaics (BPVs).** (**a**) Schematic diagram of the battery chamber. (**b**) The H-shaped dual-chamber BPV is constructed by connecting two 50 mL chambers with 5 cm diameter channels. The anode and cathode were carbon cloth (diameter of 5.0 cm). The cathode chamber is shielded from light. The anodic and cathodic chambers are separated with the proton exchange membrane Nafion 177. The anode and cathode are connected to a 1000 Ω resistor in parallel with a multi-meter to record the output voltage. All BPV experiments are operated in the static incubator at a temperature of 26°C. (**c**) The equivalent circuit of the double chamber *Dunaliella* BPVs. $R_{ct}$ is the charge transfer resistance of the cathode reaction.

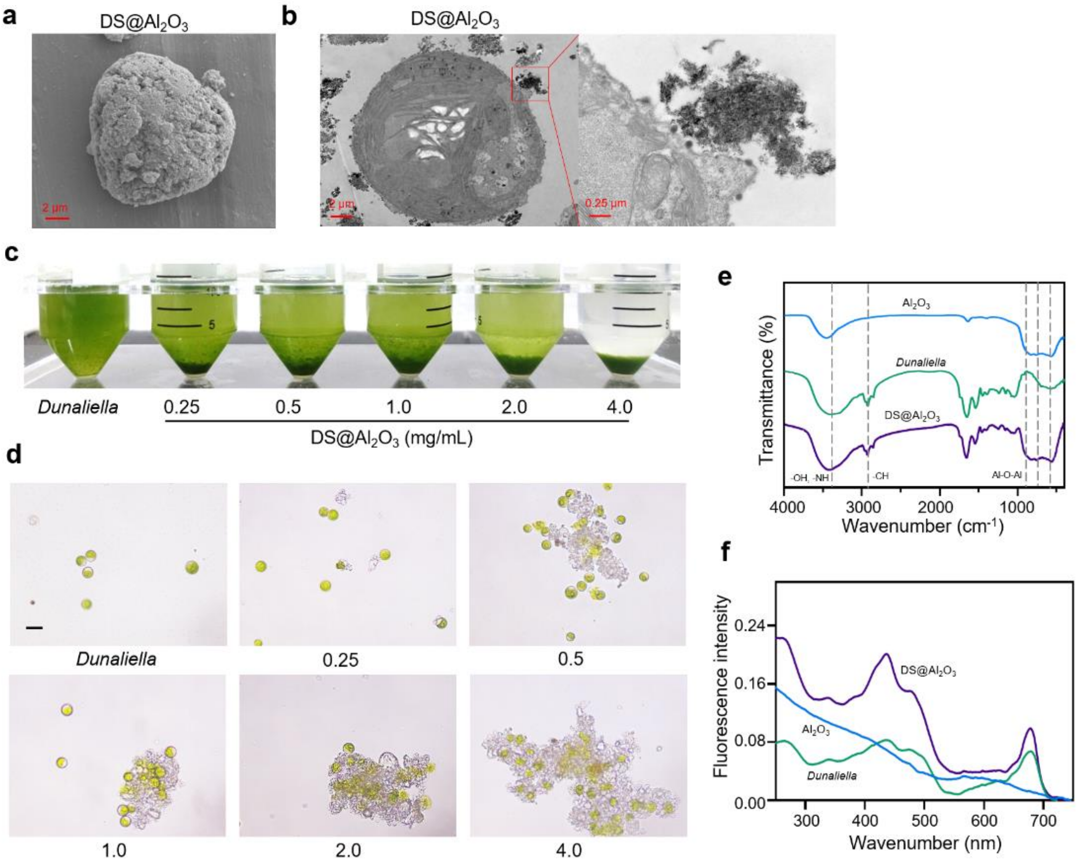


**Supplementary Fig. S6. Characterization and analysis of $Al_2O_3$-coated *Dunaliella* cells. a.** Scanning electron microscope images of *Dunaliella* cells coated with $Al_2O_3$ under scanning electron microscope. Scale bar = 2 µm. **b.** Ultrathin section images of DS@$Al_2O_3$ cells under transmission electron microscope TEM. Scale bar = 2 µm and 0.25 µm (Inset). Algal fluid (**c**) and optical microscope (**d**) of *Dunaliella* cells coated with $Fe_3O_4$ at different concentrations (0, 0.25, 0.5, 1, 2, 4 mg/mL). Scale bar = 10 µm. **e.** Infrared spectra of $Al_2O_3$, native and $Al_2O_3$-coated *Dunaliella* cells. **f**. UV-vis absorption spectrum of *Dunaliella* cell before and after $Al_2O_3$ coating.

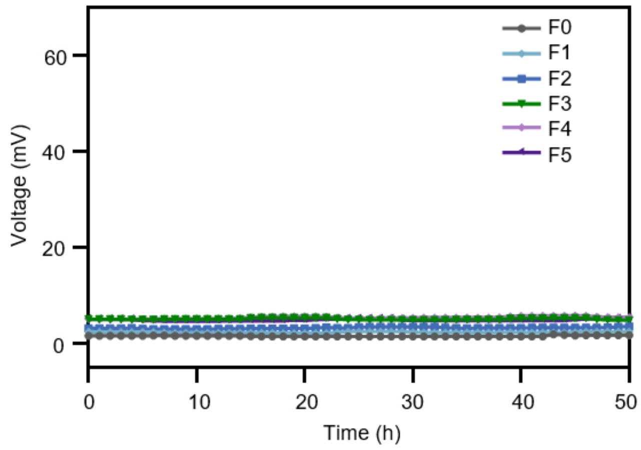


**Supplementary Fig. S7.** The controls ($Fe_3O_4$) for different $Fe_3O_4$-coated concentrations *Dunaliella* BPVs versus time curves for long-term stability and repeated cycling tests (F0: 0 mg/mL, F1: 0.25 mg/mL, F2: 0.5 mg/mL, F3: 1.0 mg/mL, F4: 2 mg/mL, F5: 4 mg/mL).

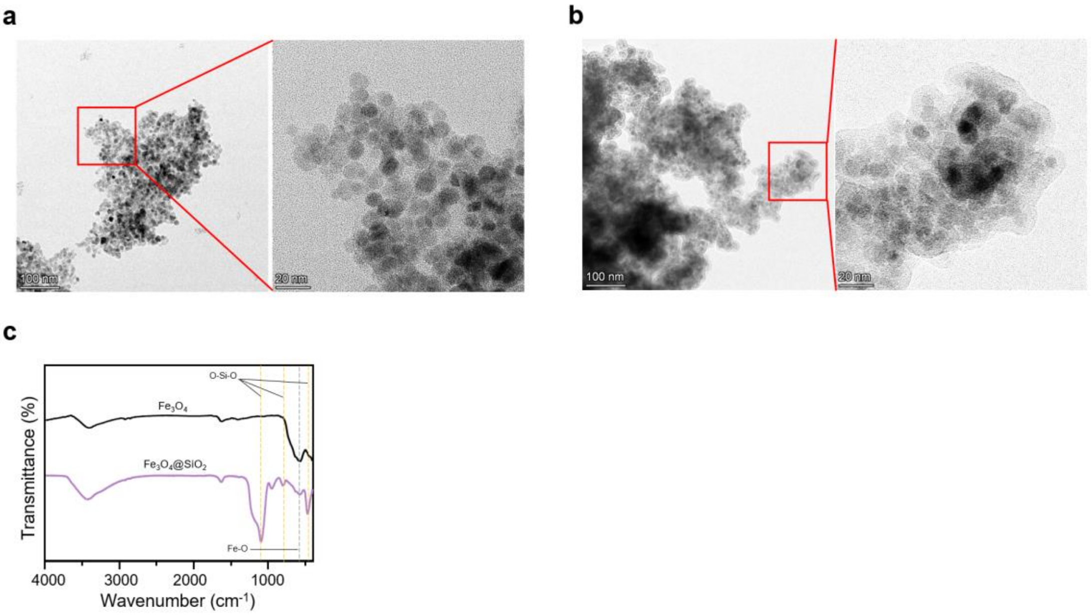


**Supplementary Fig. S8.** Transmission electron microscope (TEM) diagram of $Fe_3O_4$ (**a**) and $Fe_3O_4$@$SiO_2$ (**b**) nanoparticles. Scale bar = 100nm and 20nm (Insert) **c**. Infrared spectra of $Fe_3O_4$ and $Fe_3O_4$@$SiO_2$ nanoparticles.

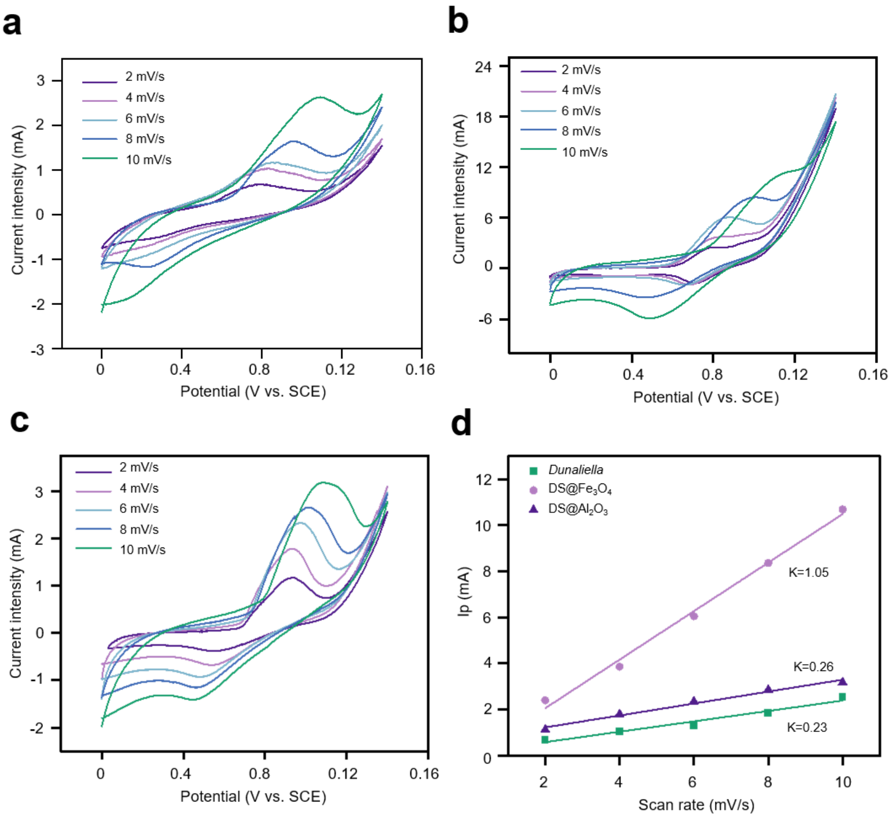


**Supplementary Fig. S9.** Cyclic voltammetry curves of native *Dunaliella* (**a**), DS@$Fe_3O_4$ (**b**) and DS@$Al_2O_3$ (**c**) BPVs and the relationship between peak current (**d**) at different scanning rate (2, 4, 6, 8, 10 mV/s).
